# Binding specificity of ASHH2 CW-domain towards H3K4me1 ligand is coupled to its structural stability through its α1-helix

**DOI:** 10.1101/2021.08.12.456084

**Authors:** Maxim S. Bril’kov, Olena Dobrovolska, Øyvind Ødegård-Fougner, Øyvind Strømland, Rein Aasland, Øyvind Halskau

## Abstract

The CW-domain binds to histone-tail modifications found in different protein families involved in epigenetic regulation and chromatin remodelling. CW-domains recognize the methylation state of the fourth lysine on histone 3, and could therefore be viewed as a reader of epigentic information. The specificity towards different methylation states such as me1, me2 or me3 depends on the particular CW subtype. For example, the CW domain of ASHH2-methyltransferase binds preferentially to H3K4me1, MORC3 binds to both H3K4me2 and me3 modifications, while ZCWPW1 is more specific to H3K4me3. The structural basis for these preferential bindings are not well understood, and recent research suggests that a more complete picture will emerge if dynamical and energetic assessments are included in the analysis of interactions. This study uses fold assessment by NMR in combination with mutagenesis, ITC affinity measurements and thermal denaturation studies to investigate possible couplings between ASHH2 CW selectivity towards H3K4me1 and the stabilization of the domain and loops implicated in binding. Key elements of the binding site – the two tryptophans and the α1-helix form and maintain the binding pocket were perturbed by mutagenesis and investigated. Results show that α1-helix maintains the overall stability of the fold via the I915 and L919 residues, and that correct binding consolidates the loops designated η1, η3, as well as the C-terminal. This consolidation is incomplete for H3K4me3 binding to CW, which experiences a decrease in overall thermal stability upon binding. Moreover, loop-mutations not directly involved in the binding site nonetheless affect the equillibrium positions of key residues.

## Introduction

Regulation of gene expression and activity at the chromatin level relies on proteins that “read”, “write” or “erase” post-translational modifications (PTMs) on histone tails and DNA. Proteins containing such domains are involved in genome organization and regulation by recognizing and modifying the PTM state of the genome compartments [1–3]. A key feature of histone tail recognition domains is that they can both recognize the general PTM state of their ligand while still have the ability to differentiate between them [4–6]. The CW domain family is a histone-tail methylation-state reader shared among numerous organisms (vertebrates, vertebrate-infecting parasites and higher plants). The name of the domain comes from its four conserved cysteines and three conserved tryptophans (Fig. 1A). The conserved cysteines coordinate a Zn^2+^ ion essential for folding, and two of the three tryptophan residues form a π-cation-based binding pocket with high affinity towards methylated lysine residues found on histone tails. The final tryptophan forms part of the hydrophobic core of the domain [7–9]. The CW domain appears within a multidomain functional context that varies from protein family to protein family [7]. One example of this is the MORC family of ATPase chromatin remodelers. Here, the CW domain recruits the proteins to the chromatin by recognizing H3K4me2/3 modifications. CW also regulates the ATPase activity of MORC3 by suspending its autoinhibition after binding to methylated H3 histone tails [10–12]. Intrestingly, the MORC4 protein, while having hight structural and sequence similarity to MORC3, is not a autoinhibiting enzyme. For MORC4 activation, CW needs to interact with DNA in addition to binding H3K4me3 [13]. Another example are the ZCWPW1 and ZCWPW2 PWWP-domain-containing proteins. Within these proteins, the CW function is unclear, but they recognize H4K20 methylation marks in addition to the H3K4me3 specificity conferred by the CW domain [8,14,15]. In the transcriptional corepressor LSD2/AOF1/ KDM1B that demethylates mono- and dimethyl H3K4 marks, CW appears to be inactive due to steric inaccessibility. However, the CW contributes to the overall structural stability of the protein and regulates the enzyme’s activity and association with mitotic chromosomes [15–18]. In the MBD protein family, the ZmMBD101 protein maintains the repressed state of the *Mutator* genes, protecting plant genomes from mutagenesis caused by transposons. The role of the CW domain in this context is still unclear [19]. CW also appears in ASHH2 (other names are SDG8 and EFS), a methyltransferase found in the small, flowering plant *Arabidopsis thaliana*. ASHH2 is involved in the regulation of gene expression by the histone H3 trimethylation at Lys-36 (H3K36me3). Within ASHH2, the CW domain preferrentially binds the H3K4me1 mark and presumably helps docking the catalytic SET domain correctly onto the histone [9,20–22].

**Figure 1:**
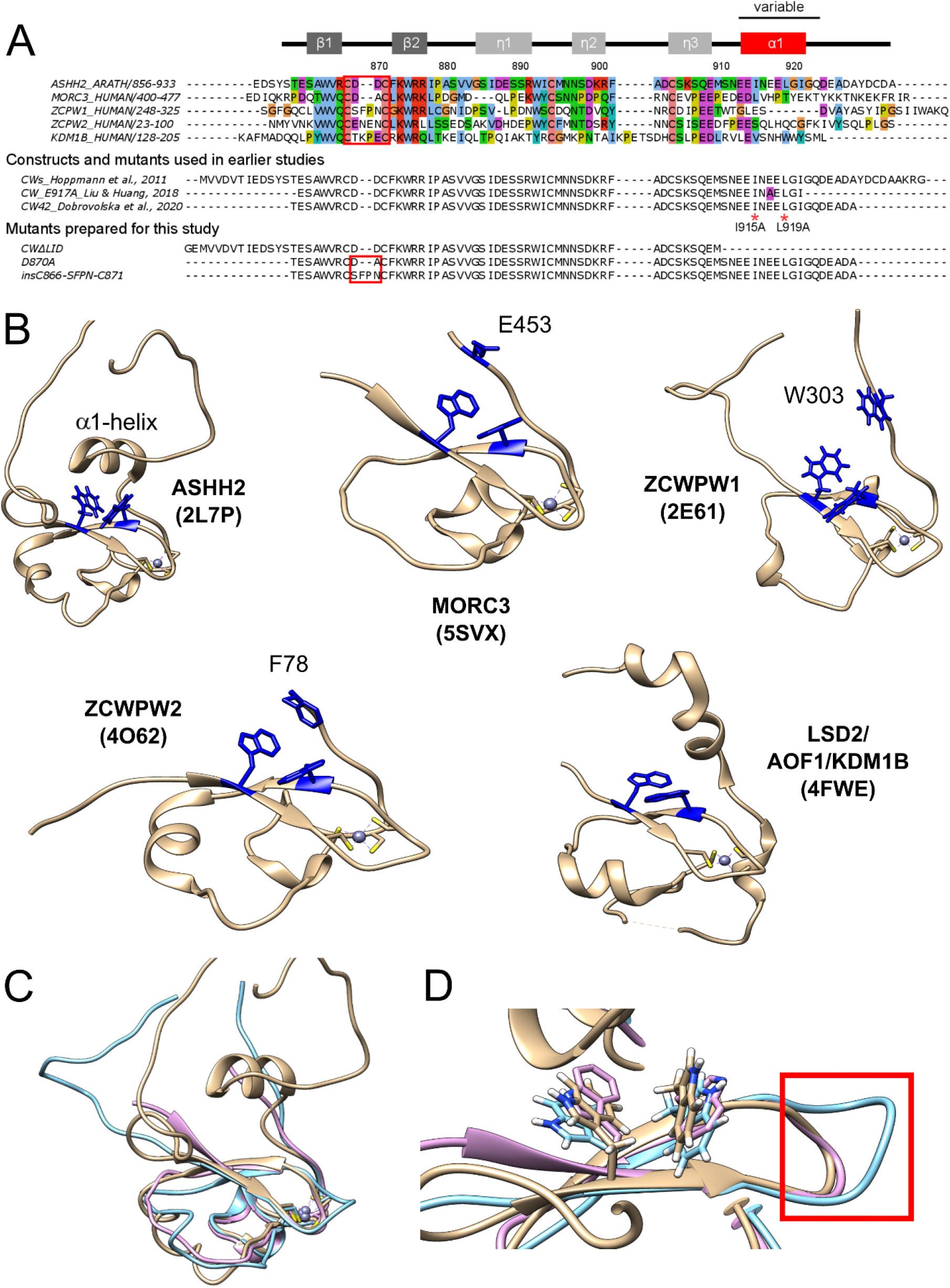
Overview of CW domains and structural analysis. A – TOP: Sequence alignment of CW-domains of ASHH2 methyltransferase (*Arabidopsis thaliana*, Q2LAE1), MORC3 protein (*Homo sapiens*, Q14149), ZCWPW1 protein (*Homo sapiens*, Q9H0M4), ZCWPW2 protein (*Homo sapiens*, Q504Y3) and KDM1B histone demethylase (*Homo sapiens*, Q08EI0). The alignment was prepared using Jalview software and UniProt entries with ClustalO default parameters, and Clustalx coloring scheme was used. MIDDLE: CW domain constructs used in earlier studies on the ASHH2 CW domain by Hoppmann *et al*., 2011, Liu and Huang, 2018 and Dobrovolska *et al*., 2020 [9,23,24]. The E917A mutation used by Liu and Huang, 2018 is marked by purple color. I915A and L919A mutations analyzed by Liu and Huang, 2018 and in this work are marked by red * symbols. BOTTOM: Sequences of the additional mutants (CWΔLID, D870A and insC866-SFPN-C871) prepared exclusively for this work. The secondary structure of CW is indicated at the top of the panel. The red squares indicate the variale loop situated between the β-strands that forms part of the binding site. These constructs also have a N-terminal sequence (“GSRRASVGSEF”), that is not shown in the figure; B – Overview of CW domain structures: ASHH2 (2L7P), MORC3 (5SXV), ZCWPW1 (2E61), ZCWPW2 (4O62) and LSD2/AOF1/KDM1B (4FWE) (tryptophans forming binding pockets and the variable C-terminal region residues, implicated in binding, are highlighted in blue); C – superposition of CW domain structures: ASHH2 (brown), MORC3 (pink), and ZCWPW1 (blue); D – close-up view of binding pocket. The loops between the β-strands subject to the insC866-SFPN-C871 mutationare indicated by red square. Graphics were prepared using the UCSF Chimera software and the pdb files represents domains in their unbound state.

The CW domain paralogs have different affinities towards different methylation states of H3K4. While the ASHH2 CW domain has higher affinity towards H3K4me1 [9,23], the rest of the known CW domains bind stronger to H3K4me2 and H3K4me3 modifications [8,10,11,15]. Earlier, factors determining the ligand methylation state specificity of CW domains have been presented and discussed [9,23,24]. The C-terminal regions of the CW domain due to its variability between paralogs were suggested to be involved in ligand specificity. For example, the CW-domain of ZCWPW1 is unique in that it has a non-conserved tryptophan residue (Trp303) at its C-terminal end (Fig. 1B), and this tryptophan finalizes the binding pocket upon binding. Mutation of this tryptophan leads to reduced affinity [8]. Its homolog ZCWPW2 has a phenylalanine residue (Phe78) in this position which also completes the binding pocket and might contribute to selectivity of the methylation state (Fig. 1B) [15]. The CW domain of MORC3 proteins, in contrast, has Glu453 residue (Fig. 1B), which finalizes the binding pocket and facilitates binding to methylated H3K4 peptides. Despite having the highest affinity towards H3K4me3 modification, its ability to differentiate between methylation states is reduced relative to what is observed for other CW domains [10,11,15]. The ASHH2-subtype possesses a unique C-terminal α1-helix located right above the tryptophan binding pocket (Fig. 1B). Although not conserved, this helix is also part of the binding pocket [23,24], as removal of this element leads to loss of ligand binding ability [9].

Recent X-ray (Liu *et al*., PDB code: **5YVX**)) and NMR (Dobrovolska *et al*., PDB code: **6QXZ**) structural studies described the CW of ASHH2 in complex with H3K4me1. The published structures agreed on the core of the complex; however, the structural data related to the α1-helix and the following C-terminal regions differ [23,24]. The studies also concluded differently with respect to binding mechanisms and determinants. The key residues for binding proposed by Liu *et al*. were L915, N916 and I919, residing on the α1-helix. These were proposed based on the crystal structure of the CW-H3K4me1 complex, where the domain construct ended right after the α1-helix (residue I921) and contained a mutation necessary for crystallization (E917A) close to the residues they identify as crucial for binding (Fig. 1A). Dobrovolska *et al*. argued that the N916 residue is not a part of the interaction mechanism, as the NMR structure of the complex showed that this residue does not make any contacts with the ligand. The studies agreed, however, that L915 and I919 locked the methylated lysine of the ligand inside the binding pocket. The construct used by Dobrovolska *et al*. to solve the structure was longer at the C-terminal than the one used in the Liu *et al*. study, which allowed elucidation of the role of the I921-Q923 region in binding. Analysis of the domain’s dynamics and flexibility using MD simulations and NMR further indicated that the binding mode of CW is described best by a conformational selection model [24]. A conformational selection mechanism requires that the protein has a fluctuating structural ensemble that includes the conformation(s) required for binding, even in the absence of ligand [25,26]. In such scenarios, any point mutations could potentially perturb the domain’s fluctuating fold, distorting the interaction mechanism that depends on fine-tuned equilibriums.

The objective of the work was to investigate the possible coupling between selectivity determinants and protein fold stability that allows ASHH2 CW-domain to preferentially recognize H3K4me1. In order to achieve this goal, we prepared two mutants affecting the positioning of the tryptophans of the binding site and two mutants involved in positioning the α1-helix in the ligand binding site. To assess the fundamental importance of this helix, we also prepared a deletion mutant, removing it entirely. We determined the affinities of H3K4me1/2/3 interacting with these mutants using ITC, performed thermal stability studies in the presence and absence of histone H3 mimicking peptides, and assessed changes to their fold using NMR fingerprinting.

## Results

### ASHH2 CW-domain interaction with H3 mimicking peptides

#### Fold assessment and chemical shift perturbation analysis

The ASHH2 CW-domain binds H3K4me1, H3K4me2 and H3K4me3, with the highest affinity towards H3K4me1 [9]. It is not known, however, whether the binding of these peptides is essentially the same or whether they affect the CW’s fold differently. ^1^H-^15^N HSQC NMR spectra of CW in complex with mono-, di-, and trimethylated ligand were acquired and compared to the unbound form. Using the backbone chemical shift assignments available for the four situations [27], the chemical shift perturbation (CSP) was calculated, and the residues involved in the corresponding complex formation were determined (Fig. 2A). The NMR data in the case of all the three histone-mimicking peptides suggests that binding causes fairly extensive structural changes that are not limited to W865 and W874 of the binding pocket. However, the HSQC spectra confirms that the CW-domain remains folded when complexed with each of the peptides (Fig. S1). This observation is in line with the flexible nature of ASHH2 CW and the proposed mechanism of conformational selection [24]. The most notable changes in chemical shifts are found in the first β-strand, perhaps related to the nascent β-sheet augmentation discussed in [24], in the conserved W891 that is not part of the binding pocket, in the α1-helix and the η1-loop (Fig. 2A).

**Figure 2:**
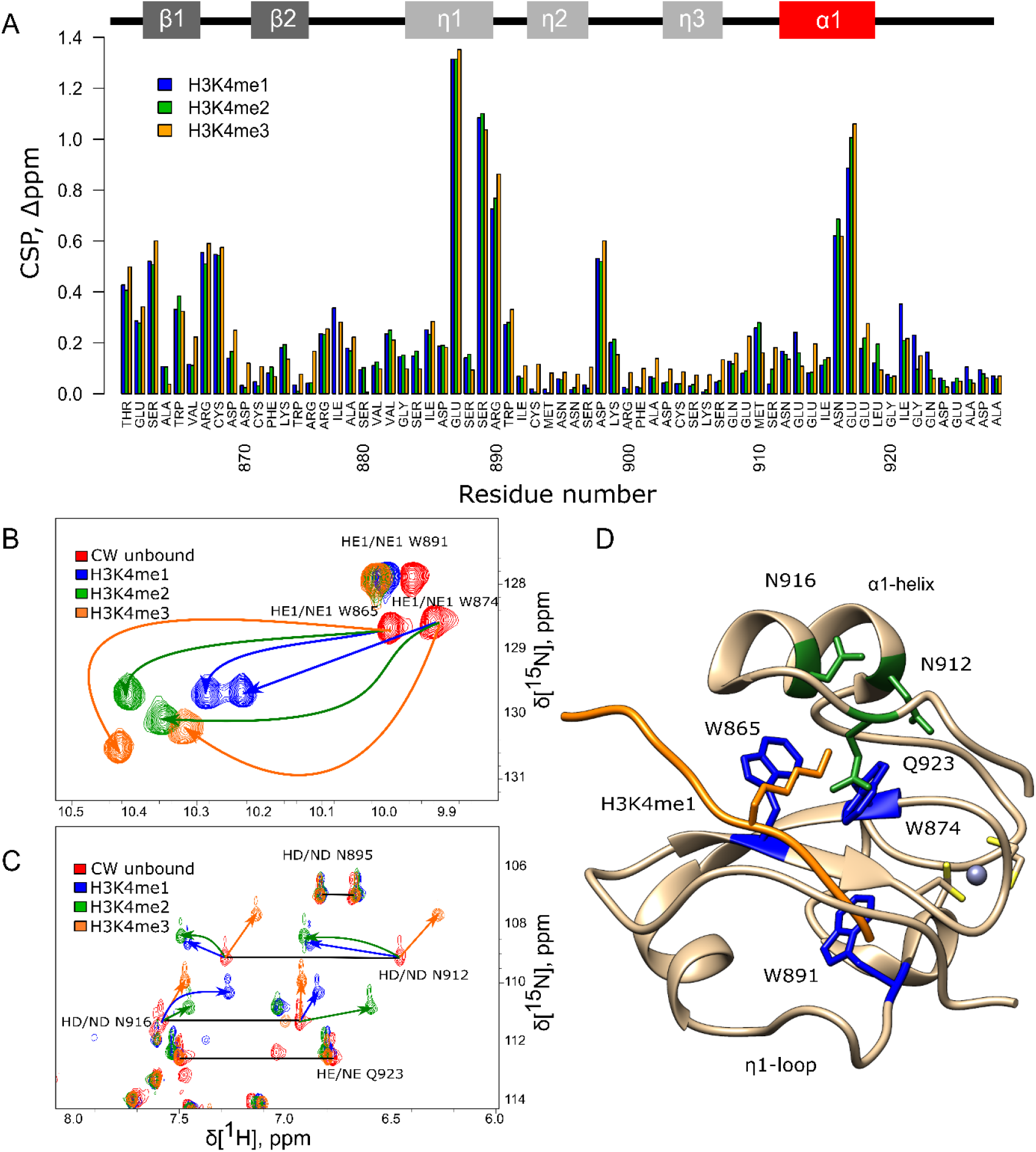
NMR analysis of CW interacting with histone mimicking peptides. A – chemical shift perturbation, calculated for CW bound to corresponding peptides (H3K4me1 blue, H3K4me2 green and H3K4me3 orange). Secondary structure of CW is indicated at the top of the panel; B and C – chemical shift of tryptophan, asparagine and glutamine side chains signals under binding of H3K4me1 (blue), H3K4me2 (green) and H3K4me3 (orange) peptides (red color – CW domain in unbound state); horizontal black lines connect signal pairs for ^15^Nε and ^15^Nδ signals of glutamines and aspargines respectively; D – side chains tryptophans (highlighted in blue), asparagines and glutamine (highlighted in green) mapped on CW structure in bound to H3K4me1 peptide (orange color) conformation (PDB: **6QXZ**); grey sphere is Zn^2+^ ion coordinated by cysteins (yellow).

The CSP data for the H3K4me1 peptide shows that E913 in the α1-helix and I921-Q923 in the C-terminal tail are involved in the binding. Earlier, it was shown that I921 and Q923 are two of the key residues establishing contacts with H3K4me1 and forming part of the final cage-like configuration around K4me1 [24]. In the case of the interaction with H3K4me2 and H3K4me3, the CSP values for these residues are less prominent, suggesting that they might not only be involved in mediating the interaction with the ligand but also contribute to selectivity towards H3K4me1. Binding to the H3K4me2 peptide has its most significant effect on W865 in the binding pocket, L873, the η1-region, M910, N916 and L919 of the α1-helix. Binding to H3K4me3 affected the structure to a somewhat larger degree, as the combined chemical shift values are slightly higher for the majority of the amino acids, especially at η2 and η3-loops (Fig. 2A). Yet, overall, the CSP data do not show much difference in how the backbone was affected by the three different peptides. The perturbation analysis takes into account only the backbone amide correlations, which may be insufficient for detection of all features of these interactions.

Amino acid side chain signals, corresponding to the NH groups of the tryptophans and asparagines and not included in the CSP analysis above, experience large shifts as a result of ligand binding. Notably, in each case of the ligand methylation state, the same NH signals had different shifts, indicating that the chemical environment around these groups was affected differently (Fig. 2B). The shifts suggest a change in the configuration of the tryptophan side-chains in the binding pocket to accommodate the additional methyl group(s) of the ligand. A similar observation for the signals corresponding to the side-chains of N912 and N916 suggests that the α1-helix also experience methylation-dependent shifts in their positions (Fig. 2C and 2D).

#### Dynamic properties of CW domain bound to histone mimicking peptides

The HSQC fingerprinting, CSP analysis and differences in the chemical shifts of the side chain of the tryptophans and asparagines together suggest there may be subtle differences in how the ASHH2 CW domain responds to different ligand methylation states. Such differences may not be readily detectable by assessing chemical shifts only, as some they reside among fluctuations in the domain’s structure and its dynamic properties that contribute little to crosspeak positions. Dobrovolska *et al*. provided a full NMR dynamics characterization for CW in the unbound and H3K4me1-bound state [24]. Here we expand part of this analysis by accumulating and comparing the hetNOE data for H3K4me1/2/3 peptides and compare them.

Overall, the average hetNOE values for the domain bound to the peptides were around 0.9, indicating moderate flexibility, except in the η1-loop, which remained quite flexible with hetNOE values as low as 0.6-0.8 (Fig. 3). Generally, the binding of the H3K4me1 peptide results in movement restriction of the unstructured C-terminal I921-Q923 region. The η3-loop residue S907 shows high flexibility, as was reported in [24], with stabilization when CW-domain is complexed with H3K4me1 and H3K4me3. Binding to the H3K4me2 peptide also had a stabilizing effect on S907, but less pronounced. When bound to the H3K4me1 peptide, a few notable outliers appear, R876, S880, A902 and C904, which displayed hetNOE values higher than 1 (Fig. 3A). R876 is located on the β2, S880 is located in the small helical feature which follows the β2 strand, and A902 is located in another helical feature, formed upon complexation with H3K4me1. The C904 residue is involved in the orientation of the Zn^2+^ ion in the core of the domain. The stability of the α1-helix region is slightly decreased for CW bound to H3K4me3 (Fig. 3C), which might have an effect on the complex stability. In sum, when complexed with me2 and me3 peptides, the dynamic behavior of CW looks more similar to the domain in its unbound state (Fig. 3B and C). This may indicate that the full shift in fold equlibrium occurs only with monomethylated ligand.

**Figure 3:**
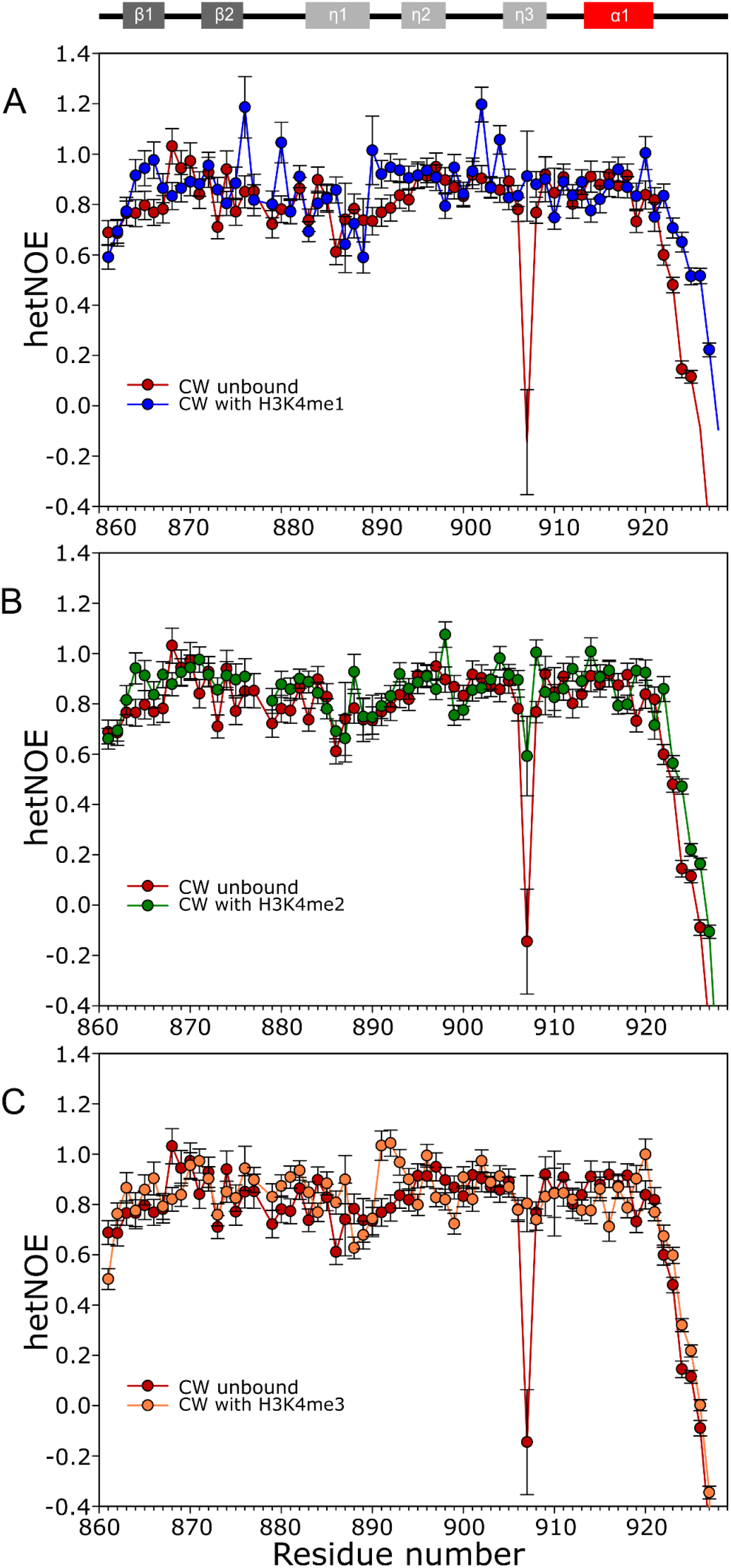
Heteronuclear ^1^H-^15^N NOEs data of CW bound to histone mimicking peptides. A – CW with H3K4me1 (blue circles, 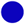); B – CW with H3K4me2 peptide (green circles, 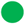); and C – CW with H3K4me3 peptide (orange circles, 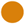). The data in each panel is overlaid with unbound CW (red circles, 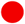). Secondary structure of CW is indicated at the top of the panels.

#### Thermodynamical characterization of the interaction

The NMR analysis suggests that CW, when complexed with the peptides, sample available conformations differently at equilibrium. If a given methylation state fail to stabilize the complex, this should be reflected in its thermal stabilities. The CW-domain in its unbound and bound state was therefore subjected to thermal denaturation analysis monitored by intrinsic tryptophan fluorescence. Complexation with H3K4me1 and H3K4me2 peptides increased the thermal stability of the domain: T_m_ = 67.3 °C without peptide, T_m_ = 71.4 °C with H3K4me1 and T_m_ = 71.6 °C with H3K4me2. In contrast, interaction with H3K4me3 reduced this stability to T_m_ = 64.9 °C (Fig. 4A and B).

**Figure 4:**
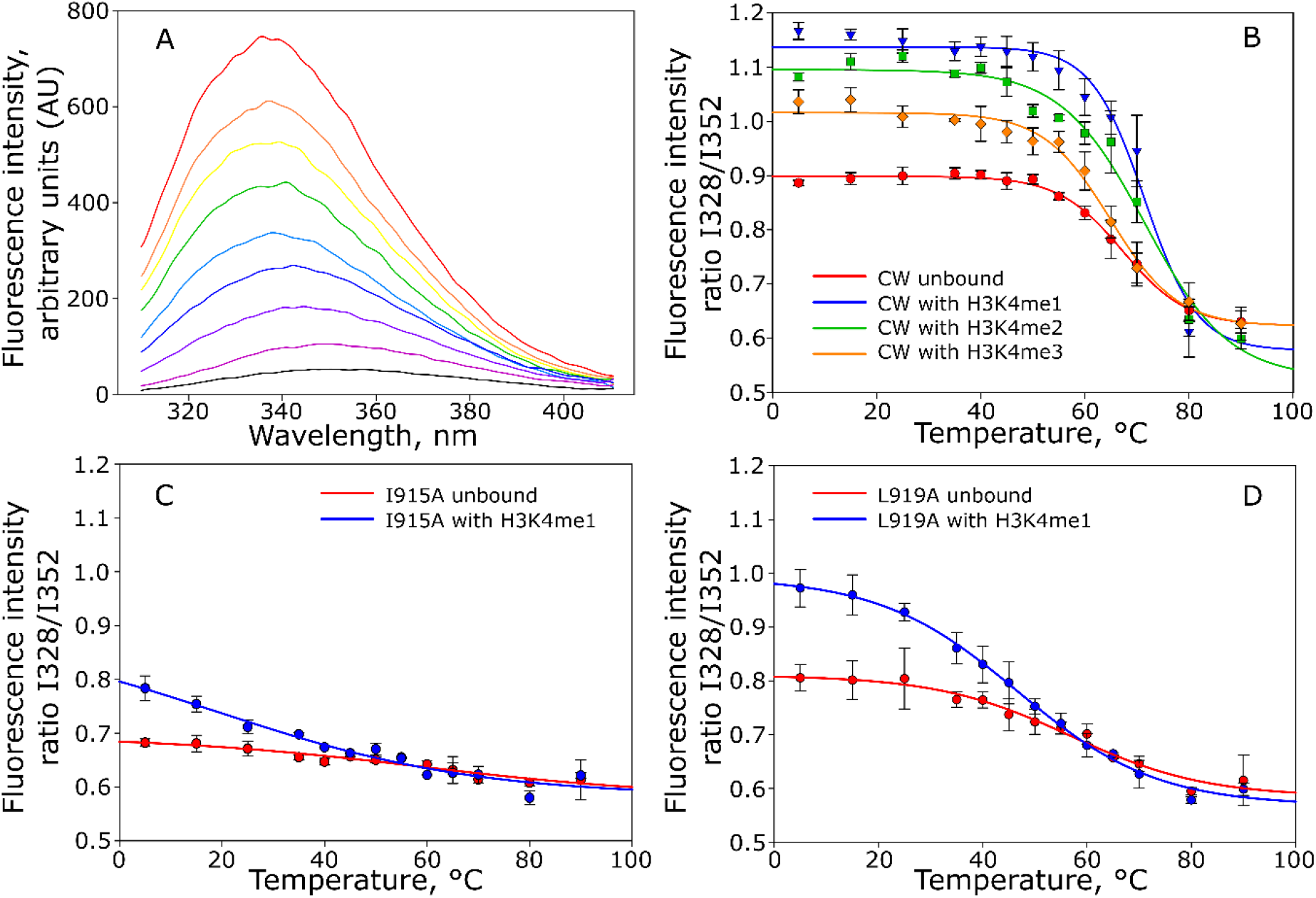
Thermal denaturation of CW monitore by intrinsic tryptophan fluorescence in unbound and bound states. A – Representative fluorescence traces for wild type CW bound to H3K4me1 (only selected traces are shown); B – melting curves for unbound CW domain 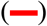 and CW domain in presence of H3K4me1 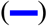, H3K4me2 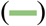 and H3K4me3 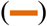 peptides; C – melting curves for unbound CW I915A mutant 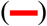 and in presence of H3K4me1 peptide 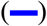; D – melting curves for unbound CW L919A mutant 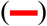 and in presence of H3K4me1 peptide 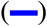. Number of replicates = 3.

In order to study thermodynamical forces underlying ligand binding and selectivity, we performed ITC measurements of CW interacting with H3K4me1/2/3 peptides. Interaction with H3K4me1 peptide resulted in K_d_ = 1.31 ± 0.32 μM, binding enthalpy ΔH = −89.25 ± 8.58 kJ/mol and entropy ΔS = −192.42 ± 15.31 J/mol.K. Interaction with H3K4me2 resulted in K_d_ = 4.61±0.28 μM, binding enthalpy to ΔH = −83.64 ± 3.11 kJ/mol and entropy to ΔS = −178.37 ± 10.82 J/mol.K. Interaction with H3K4me3 displayed weaker binding relative to both H3K4me2/3, with a dissociation constant K_d_ = 14.18 ± 0.74 μM. Corresponding binding enthalpy and entropy were −60.14 ± 1.38 kJ/ mol and −108.90 ± 5.00 J/mol.K, respectively. Stoichiometric coefficients were in the range of 0.82 - 1.08. As a control, the unmodified H3K4 peptide was used and showed no binding. Results of the analysis are summarized in Fig. 5 and Table S3, representative isotherms are shown in supplementary Fig. S2.

**Figure 5:**
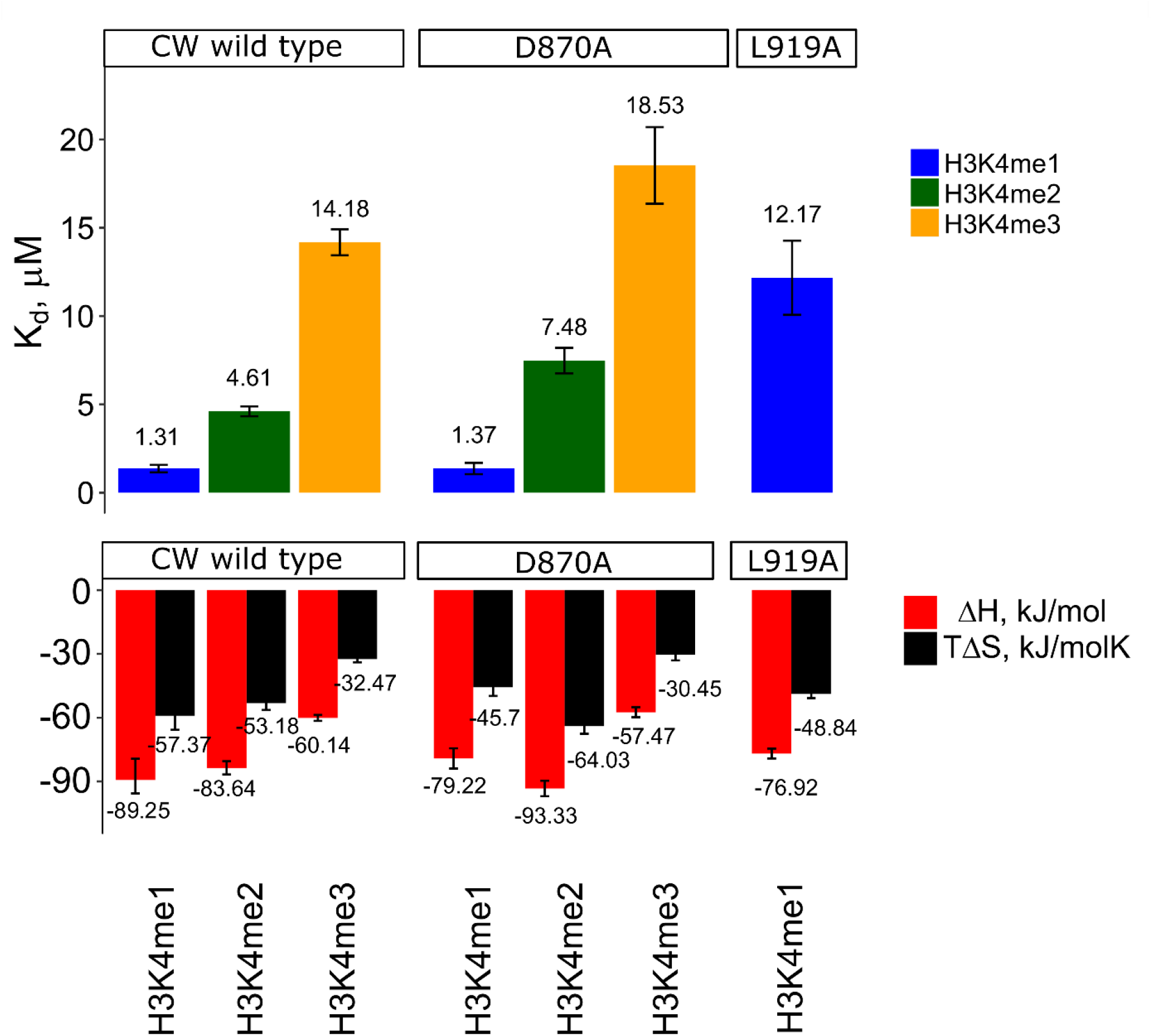
K_d_, ΔH and TΔS values determined by ITC. Data shown only for WT CW and mutants D870A and L919A, as insC866-SFPN-C871 and I915A did not produced interpretable isotherms. Number of replicates = 3. Error bars show standard deviation. Representable isotherms can be found in supplementary Fig. S2.

### Effect of α1-helix mutations on binding affinity, domain conformation and thermal stability

A detailed structural study of CW binding to H3K4me1 showed that the α1-helix and the disordered region following it contribute critically to binding. A group of CW residues residing in this region establish direct NOE contacts with the ligand and comprise L919, I915 and Q923 [24]. Hoppmann *et al*. showed that without this part, the protein is no longer capable of binding a ligand. It is, however, unclear whether this is due to loss of fold, or inability to retain the ligand in the otherwise intact binding site. To assess the impact α1-helix has on the fold of the protein, the CWΔLID mutant (residues 847-910 removed) was generated, and a ^1^H NMR spectrum was acquired. Compared to the wild type, the spectrum of the CWΔLID shows fewer and broader peaks spread across a narrower region, which confirms that removal of the α1-helix leads to loss of fold (Fig. 6A). This finding confirms the joint importance of both ligand binding and fold maintainance of the α1-helix and the C-terminal region.

**Figure 6:**
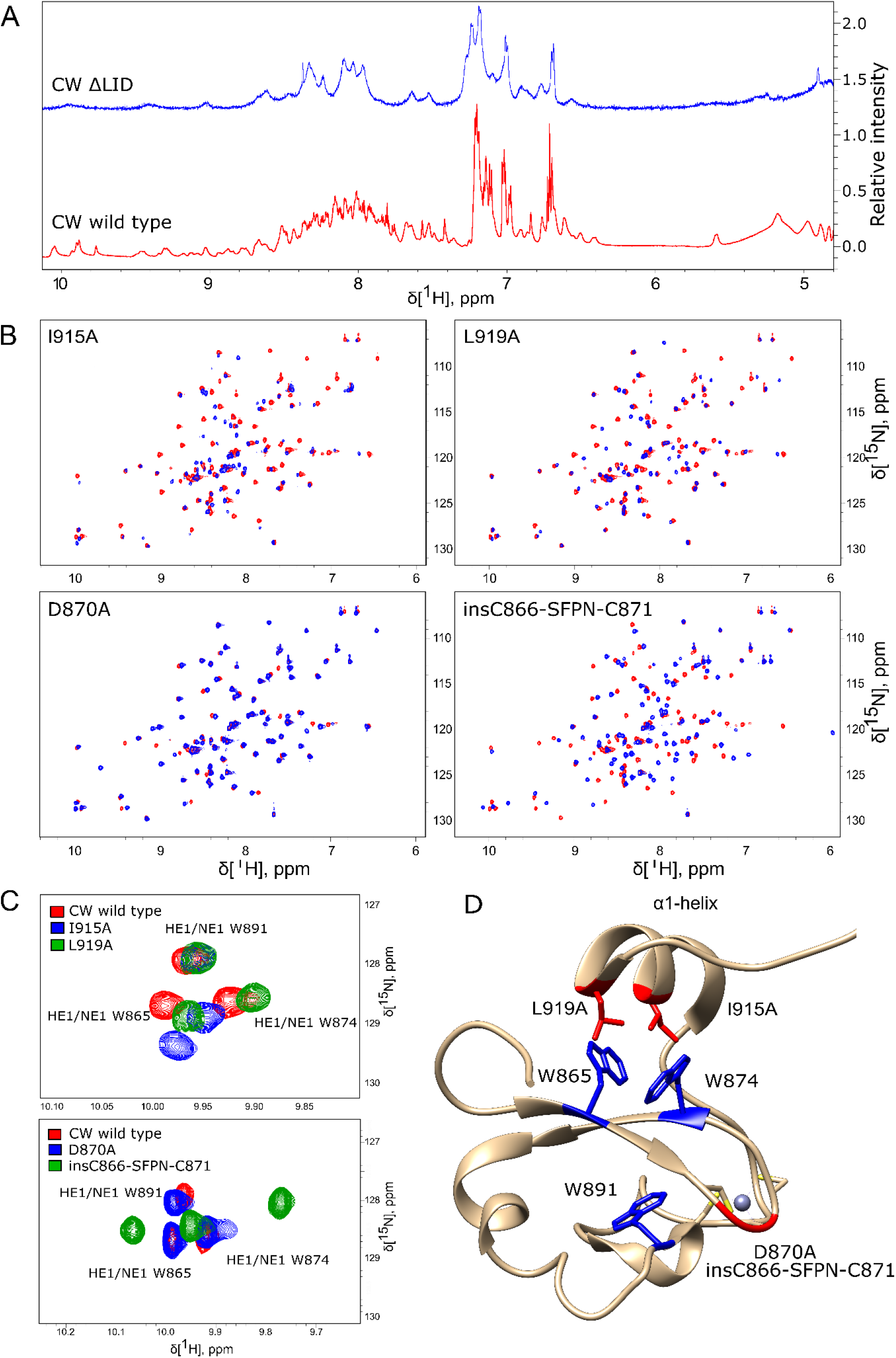
Mutation induced structural disturbances analyzed by NMR. A – proton spectrum of CW ΔLID (blue) compared to the proton spectrum of wild type CW (red). The CW ΔLID trace has been shifted up by 1.5 relative units to make comparison easier. B – ^15^N HSQC spectra of I915A, L919A, D870A and insC886-SFPN-C871 (blue color) overlaid with spectra of WT CW (red color). C – the effect of mutations in α1-helix (red – WT CW, blue – I915A, green – L919A (top panel)), and the effect of mutations in the loop between β-sheets (red – WT CW, blue – D870A, green – insC866-SFPN-C871 (bottom panel)). D – location of the point mutations (red sticks) and insertion mutations (red backbone trace) mapped on the CW structure (PDB: **2L7P**). The tryptophans are highlighted in blue; grey sphere is Zn^2+^ ion coordinated by cysteins (yellow).

To examine the role of the α1-helix in domain stability and specificity, we prepared I915A and L919A mutants. Their ^1^H-^15^N HSQC NMR spectra showed that the protein remains folded (Fig. 6D and E). Compared to the wild type, I915A and L919A mutations show animpact on the overall fingerprint, as more than 60% of the peaks were shifted substantially (more than 0.2 ppm) (Fig. 6B). Thermal denaturation experiments showed that compared to the native CW domain (T_m_ =67.3 °C), the L919A mutation significantly decreased the stability of the protein by 10.4 °C (T_m_ = 56.9 °C) (Fig. 4C and D). Moreover, the binding of H3K4me1 decreased it further, by more than 10 degrees in addition (T_m_ = 46.7 °C). This behaviour strikingly contrasts with the wild-type where the ligand increases the T_m_ by ≈ 5 °C. I915 mutation had a more dramatic impact on the fold stability, as it was not possible to fit a curve to the data from denaturation analysis, as the data does not display a sigmoidal model.

Changes in the binding preference for I915A mutant was previously evaluated using ITC by Liu *et al*., with the conclusion linking it to a change in specificity from H3K4me1 to H3K4me3. When we performed the ITC experiment with the I915A mutant using our slightly longer construct (CW42), it was not possible to obtain an accurate K_d_ and other thermodynamic parameters, presumably due to its weak interaction with the peptides. The interaction of L919A mutant with H3K4me2 and H3K4me3 peptides could not be characterized either. In the case of H3K4me1 interaction with L919A, the K_d_, the binding enthalpy, and entropy were determined to be 12.17 ± 2.10 μM, −76.92 ± 2.31 kJ/mol (ΔH), and −163.8 ± 6.61 J/mol.K (ΔS), respectively. The stoichiometric coefficients were in the range of 0.86 - 0.88. The results are summarized in Fig. 5 and Table S3. The shift in specificity reported by Liu *et al*. was not confirmed in our experiments [23].

### Binding pocket flexibility as possible determinants for binding preferences

Binding mode and ligand specificity of each CW subtype must ultimately arise from the environment provided by the binding pockets. To compare structures, we performed sequence and structural alignments of CW domains from ASHH2 (2L7P), MORC3 (5SVX), and ZCWPW1 (2E61) (Fig. 1A). Structural superimposition of ZCWPW1, ZCWPW2, MORC3 (human and mouse), and ASHH2 CW revealed different angles between the tryptophans side-chains in the binding pocket (Fig. 1D). ASHH2 has the narrowest angle (65°), which might partially explain its specificity towards H3K4me1. In the remaining structures, with a reported preference towards H3K4me3, the angle varies between 77° and 113°. Upon ligand binding, the angle increases, widening the binding pocket (results are summarized in Table 1). This expansion might be possible due to the flexibility of the loops surrounding the binding pocket, with ASHH2 having insufficient flexibility to open up the pocket for stable binding to the H3K4me3 ligand. The gap in ASHH2 and MORC3 sequence alignments (Fig. 1A) translates into a shorter loop between two β-sheets, compared to ZCWPW1 (Fig. 1D). Inside this loop, MORC3 has alanine in the place of ASHH2’s D870, and ZCWPW1 has the sequence S867-FP-N870 inserted relative to ASHH2. These variations in length and composition of the loop may impact the orientation of the β-strands and thus on also the positioning of the conserved tryptophans in the binding pocket.

**Table 1.** Angles between tryptophan side chains forming the binding pocket. Tryptophan angle measurements were performed manually with Screen Scales from Talon-Designs LLP and automatically by fitting a plane through the coordinates of the atoms in the two tryptophan side chains and solving the angle between the two planes.

| Uniprot ID | PDB ID | Ligand | Angle,<br>manual | Angle,<br>automatic | Highest<br>specificity | Reference |
| --- | --- | --- | --- | --- | --- | --- |
| ASHH2_ARATH | 2L7P | Unbound | 65.9 | 64.9 | H3K4me1 | [9,23,24] |
|  | 6QXZ | H3K4me1 | 94.4 | 94.3 |  |  |
|  | 5YVX | H3K4me1 | 113.0 | 112.6 |  |  |
| MORC3_HUMAN | 4QQ4 | H3K4 | 99.6 | 99.7 | H3K4me3 | [10,15] |
|  | 5SVY | H3K4me1 | 99.1 | 98.1 |  |  |
|  | 5SVX | H3K4me3 | 100.5 | 100.1 |  |  |
| MORC3_MOUSE | 5IX2 | H3K4 | 78.0 | 76.8 | H3K4me3 | [11] |
|  | 5IX1 | H3K4me3 | 89.9 | 88.6 |  |  |
| ZCWPW1_HUMAN | 2E61 | Unbound | 104.1 | 104.3 | H3K4me3 | [8] |
|  | 2RR4 | H3K4me3 | 107.6 | 105.7 |  |  |
| ZCWPW2_HUMAN | 4O62 | H3K4me3 | 113.2 | 111.2 | H3K4me3 | [15] |

Loops have been implicated in the binding mechanism of the CW domain, where they regulate the positioning of α1-helix (η3-loop and the post-helix C-terminal coil). Additionally, the η1-loop also interact with the ligand upon complex formation, and together these unstructured elements mediate binding and stabilize the complex [24]. To study the role of the loop that connects the two β-sheets scaffolding the binding pocket, the D870A and insC866-SFPN-C871 mutants were prepared, with mutations that correspond to the composition of MORC3 and ZCWPW1 loops respectivelly. The folded states of the mutants were verified by ^15^N HSQC fingerprinting (Fig. 6B). The D870A mutant had little effect on the structure, affecting only around 7% of the signals in the fingerprint. The insC866-SFPN-C871 insertion mutant, had a more pronounced effect on the structure, resulting in a shift of around 45% of signals (Fig. 6B). Changes in the binding preferences of the D870A and insC866-SFPN-C871 mutants were evaluated by ITC.

It was not possible to obtain K_d_ and other thermodynamic parameters for the insC866-SFPN-C871 mutant due to weak interactions with all three histone mimicking peptides. Change in selectivity was not observed, and interaction of the D870A sample with H3K4me1 was comparable to WT values (K_d_ = 1.37 ± 0.32 μM, binding enthalpy ΔH = −79.22 ± 4.81 kJ/mol and entropy ΔS = −153.27 ± 13.75 J/mol.K). This mutation gave decreased binding affinity towards H3K4me2 and me3 peptides, however: interaction with H3K4me2 resulted in K_d_ = 7.48 ± 0.72 μM, ΔH = −93.33 ± 3.65 kJ/mol and ΔS = −214.77 ± 12.06 J/mol.K. For H3K4me3, these binding parameters were K_d_ = 18.53 ± 2.17 μM, ΔH = −57.47 ± 2.37 kJ/mol, and ΔS = −102.14 ± 8.90 J/mol.K. The stoichiometric binding coefficients were in the range of 0.96 - 1.01. Results are summarized in Fig. 5 and Table S3 with representative isotherms shown in supplementary Fig. S2.

### Effect of mutations on the environment of the tryptophan side chains

Chemical shift analysis of the unbound state and of the H3K4me1-3 bound situations suggested that side chains were more sensitive to the binding pocket environment compared to their respective backbone atoms. Therefore, we also compared the spectra of insC866-SFPN-C871, D870A, and I915A, L919A to the spectra of the WT with a focus on sidechain NH crosspeaks. Signals from the ε-NH nuclei of the aromatic side chain of tryptophans are located in the ^1^H 7-11 ppm and ^15^N 128-130 ppm region of ^1^H^15^N HSQC spectra. Mutations in the α1-helix did not affect the core tryptophan W891 but significantly affected the tryptophan side chains in the binding pocket (W865 and W874). For L919A, ^15^Nε resonances were shifted upfield, maintaining a similar pattern as in the wild type domain. The I915A mutation affected the side chains somewhat differently, with the ^15^Nε resonance shifted downfield in the case of W865 (Fig. 6C). The mutant with the MORC3-like loop maintained a chemical environment similar to the WT situation, with slight shift of the signal from the core tryptophan side chain (W891). The mutant with the ZCWPW1-like loop had the most pronounced effect on the core tryptophan (W891), and also shifting the ^15^Nε resonances of binding pocket’s tryptophan side chains upfield (W865 and W874) (Fig. 6C).

## Discussion

Analysis and comparison of known of CW domains structures suggested that the specificity of the domain might be related to the geometry of the binding pocket, formed by tryptophan side chains, fluctuating conformers and flexible nature of the protein. Exploring the dynamic and energetic determinants of the ASHH2 CW-domain and several mutants, we found that the only in the case interaction with H3K4me1 did binding result in a fold-stabilization. We also found that the α1-helix was a requirement for the domain fold, and that mutations in loop structures had a marked effect on it thermal stability, fold and ability to interact with H3K4me1/2/3 histone mimicks.

Early characterization of ASHH2 CW-domain behaviour was reported by Hoppmann *et al*., and expanded with the structures of the bound state by Liu *et al*. and Dobrovolska *et al*. In this study, the K_d_ values for CW interaction obtained using ITC do not match entirely with the values obtained by Hoppmann *et al*. for H3K4me2 (K_d_ of 2.1 μM vs. 4.61 μM in the present study) and H3K4me3 (K_d_ of 4 μM vs. 14.18 μM in the present study), but are comparable in the case of H3K4me1. Comparing to Liu *et al*. data, the values reported are in better agreement. The discrepancy may be explained by differences in methods and protein constructs used. Hoppmann *et al*. employed SPR, which is sensitive to mass transfer, re-binding, and surface effects [28,29]. In addition, the study used a GST-ASHH2-CW construct, where the presence of the GST-tag could potentially affect the interaction [30]. However, both studies conclude that H3K4me1 is the strongest binder, whereas H3K4me3 is the weakest.

The ITC data shows that peptide binding is enthalpy-driven, and the difference in affinity arises from different enthalpy-entropy contributions [31,32]. H3K4me1 shows the most favourable enthalpy contribution of the three histone-mimicking peptides. Interaction with the other peptides showed a smaller enthalpy contribution. As the change in enthalpy in non-strict terms characterizes the number of non-covalent bonds formed upon complexation [33], this can be interpreted as the formation of fewer non-covalent interaction contacts for H3K4me2 and -me3 cases. Interaction with these peptides is also characterized by an increase in entropy terms when compared to H3K4me1 interaction. This is consistent with a higher degree of mobility of these complexes. The results show that the dominant driving force is the enthalpy overcoming an entropic barrier, which is consistent with the conformational selection mechanism, as in the lock-and-key model the interaction is driven by solvent gain in entropy [33–35]. Taking together these observations, chemical shift perturbation, dynamics data and results published by Dobrovolska *et al*., we argue that the flexible I921-Q923 region of CW is able to make contacts only when complexed with H3K4me1 peptide, remaining mobile in other cases, and by that it is involved in selectivity towards H3K4 modification.

Conformational analysis with NMR chemical shifts perturbation and thermal stability measurements showed that the conformation of the domain differs when interacting with different methylation states. Dynamic data showed that the structure of CW is adjusted to the correctly bound form by restricting the mobility of the flexible η1, η2, η3-loops and I921-Q923 unstructured region. Interaction with the H3K4me3 peptide perturbed the conformation to a greater extent than the lower methylation states of peptides, and reduced fold stability markedly. These observations are in agreement with the model presented by Liu *et al*., where steric hindrances between the binding pocket and trimethylated group were discussed. However, the impact of this steric hindrance is not restricted to the binding site, but appears to affect the whole fold.

Compared to other known structures of CW-domains, the ASHH2 subtype has the α1-helix, whose function seems to be related to the specificity of the domains [23]. We show that truncation of the helix eliminates binding by disruption of the domain fold. Point mutations of the amino acids whose side chains are oriented towards the tryptophan binding pocket confirmed that the structure is stabilized by these residues. Comparing the I915A and L919A mutations, we conclude that I915 contributes the most to structural stabilization and is also crucial for maintaining the fold optimal for binding. The I915A mutation had a pronounced effect on the overall structure and chemical environment around the tryptophan side chains of the binding pocket, and thus, likely affected its geometry, which resulted in loss of quantifiable interaction. It was not possible to determine the T_m_ for the I915A mutant with intrinsic tryptophan fluorescence spectroscopy, as the data could not be fitted by a sigmoidal curve. Knowing from HSQC fingerprinting that the I915A mutant remains folded, such behaviour can be explained by the exposure of tryptophans to the environment [36]. One possibility is that the I915A mutation could distort or displace the α1-helix exposing the tryptophan side chains, as also seems from the change in chemical environment around indole rings of the tryptophans in the HSQC side chain analysis. The L919A mutation showed decreased affinity to H3K4me1 arising from changes in the balance between enthalpy and entropy contributions of the interaction. Higher values for enthalpy and entropy indicate that the binding of the peptide was not complete relative to the native situation, and thermal stability measurements indicated a notable destabilization of the complex. These observations are in agreement with conclusions from Liu *et al*., where they demonstrated the importance of I915 and L919 in the formation of the binding pocket. However, their ITC results and conclusion about selectivity of the domain have to be treated with care, as their reported stoichiometric coefficient values did not approach 1:1 binding, as would be expected for a well-optimized system. This deviation might also indicate a low quality sample [37].

To explore the hypothesis that the specificity of the domain might be related to the flexibility of the binding pocket, mutants with variable inter-β-strand loop were prepared and analyzed. These mutations did not change the binding preference, but the composition and length of the loop between β-strands did have an impact on binding affinity. The CW-Z1loop mutant also had a significant effect on the fold of the domain. The CW-M3loop mutation did not affect the chemical environment of the tryptophans in the binding pocket, and as the result, the interaction of the CW-M3loop mutant with histone mimicking peptides similar to the WT. The interacation is driven by enthalpy, but with slightly reduced affinities towards H3K4me2 and H3K4me3. In the case of the CW-Z1loop mutant, the interaction was significantly weakened, and it was not possible to find the biding isotherm by ITC. This loss in affinity can be explained by a high degree of structural perturbance of the domain, as indicated by its HSQC fingerprint.

Ligand binding has different effects on the protein stability depending on the interaction system. Typically, a ligand stabilizes the structure, but in case of binding to partially unfolded intermediate states of proteins, the stability can be decreased [38–41]. Intrinsically disordered proteins interacting with ligands are characterized by the formation of secondary structure elements upon binding [42,43]. This is, in turn, associated with a reduction in dynamics of flexible loops, shrinking in protein hydrodynamic size and reduction in conformational entropy [41,44–46]. The affinity of the interaction arises from enthalpy-entropy compensations, where the most stable complex is formed with the most favorable thermodynamic terms [47]. With this in mind, our observations indicate that the unique specificity of the ASHH2 CW domain towards H3K4me1 modification is coupled with the stabilization of its fold. The stability itself is maintained by the α1-helix via the I915 and L919 residues, and increases upon complexation with the monomethylated ligand. Consolidation also involves the η1, η3-loops and C-terminal coil, and possibly also β-sheet induction. This consolidation is incomplete and therefore energetically unfavorable when the domain interacts with H3K4me2 and H3K4me3 peptides.

There is unfortunately little structural information available on how the CW domain might act in the context of the full ASSH2 protein. It is possible that selectivity towards histones are increased in the full context of the enzyme, or that the CW domain elicits changes to the domain organization of the full protein when bound. The work done by Dong *et al*. and Xu *et al*. provide some interesting findings regarding this [21,22]. These groups performed methyltransferase assays with a radiography labeled methyl group donor. Xu *et al*. showed that a truncated construct of ASHH2 lacking the CW domain and retaining only its catalytically relevant methyltransferase domains, is able to methylate *in vitro* H3 histone as well as H4. Dong *et al*. in their study used a full-length ASHH2, which was shown to “write” a methylation mark on H3 histone only. These observations together immediately suggest that the CW-domain of ASHH2 methyltransferase might function by restricting the SET domain activity specifically towards H3 histones.

## Materials and methods

### Materials

The Histone H3 tail mimicking peptides were synthetized by Lifetein (sequences are listed in Table S2) and had 95% purity (assessed by mass spectrometry). Buffer components and chemicals were purchased from Sigma-Aldrich. D_2_O, ^15^N enriched (99%) NH_4_Cl and ^13^C enriched (99%) glucose were purchased from Cambridge Isotopes Laboratories, Inc. (Tewksbury, MA, USA), and SVCP-Super-3-103.5 NMR tubes were acquired from Norell Inc. (Morganton, NC, USA).

### Analysis of known structures

To determine the angles between the tryptophan residues in different CW domains, a multiple structure alignment was performed on the A chain of known CW domain structures using the POSA web tool [48]. Results were visualized using UCSF Chimera [49] and PyMOL, and the measurements of the angles between the two tryptophan residues were measured manually with Screen Scales from Talon-Designs LLP, and automatically by fitting a plane through the coordinates of the atoms in the two tryptophan side chains and solving the angle between the two planes. Git code is available at https://github.com/oodegard/CW_domain_paper. Sequence alignment was performed using Jalview software [50] with Clustal O algorithm with default parameters.

### Protein expression and purification

Protein expression and purification was performed as described in [27]. Mutant versions of ASHH2 CW-domain were generated using site-directed mutagenesis and PCR. The primers used are listed in Table S1. After PCR, the reaction mix was treated with DpnI, and linearized DNA was ligated at room temperature for 20 minutes in reaction containing PCR product, NEB4 buffer, T4 ligase, PNK T4 Kinase, 1 mM ATP and 10 mM DTT. Mutant CW-constructs were expressed and purified as described above. All constructs were verified by sequencing.

### Isothermal titration calorimetry (ITC)

For all measurements, the temperature and stirring rate were kept constant at 25 °C and 300 rpm, respectively. Sample concentration was 50-180 μM and the enthalpy of binding was determined by stepwise titration of 400-1800 μM histone peptide. Each CW sample (wt and mutants) were analyzed with H3K4me1, H3K4me2, H3K4me3 (Peptide sequences listed in Table S2). For each experiment, 2 μL of peptide was injected 22 times with 300 seconds intervals between injections. Experiments were performed in triplicates. Both protein and peptide were dissolved in T7 buffer (25 mM Tris-HCL pH 7.0, 150 mM NaCl, 1mM TCEP) and the heat of peptide dilution into T7 buffer was subtracted from the measurement. Binding parameters were determined from the integrated peak areas with independent modelling, using the TA Instruments NanoAnalyze V 2.4.1 software.

### Thermal denaturation

The melting point of the different CW domain constructs, defined as the temperature at the inflection point of I328/I352 during heat denaturation, was determined with and without ligand. Each sample was subjected to a temperature gradient ranging from 5 to 90 °C. Each 5 °C – 10 °C increase in temperature was allowed to come to thermal equilibrium before measurements were taken. Fluorescence was recorded in the range between 310-410 nm with a scanning rate of 100 nm/min, 2 scans per sample. Each sample was analyzed in triplicates. The cuvette was equipped with a lid to prevent evaporation from the sample. Samples were dissolved in T7 buffer, protein concentration was 10-20 μM and ratio of protein to peptide was 1:6. Corresponding blanks (buffer or peptide solution) were subtracted from the measurements. Ratio of fluorescence intensities at 328 nm and 352 nm were plotted against temperature. A sigmoidal curve with four variables (Equation 1) was fitted between parallels to obtain the inflection point (taken as the melting temperature, T_m_). Curve fitting was performed with SigmaPlot v13.0 followed by one-way ANOVA statistical test with p-value threshold of 0.05.

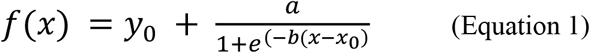

Here, *f(x)* is a function of temperature, *y*_0_ is the slope of the pre-transition state, *a* is the slope of post-transition state, *x*_0_ is the infection point of the curve, and *b* is the slope of the transition state at the inflection point *x*_0_.

### NMR spectroscopy

Protein samples were dissolved in NMR buffer (20 mM Phosphate buffer pH 6.4, 50 mM NaCl, 1 mM DTT) with 10% D_2_O. All NMR data was acquired on a Bruker Ascend 850 MHz instrument, fitted with a cryogenically cooled TCI probe. Data was processed in TopSpin 3.5 pl 6.2. ^1^H^15^N-HSQC fingerprints were acquired using the f3 channel, ^15^N decoupling during acquisition, water flip-back pulse and echo/anti-echo-TPPI gradient selection [51]. Typical parameters were: p1 7.06 μs, d1 1.0 s, SW 15.92 ppm (F2) and 35 ppm (F1), o1p 4.7 ppm, o3p 35 ppm. TD was set to 2048 and 128 in F2 and F1, respectively. Data was acquired using 50% non-uniform sampling. For processing, a forward complex linear prediction was applied in F1 up to 256 points and zero-filled up to 512 points. The FIDs were apodized using a squared cosine function in both dimensions prior to Fourier-transformation. HNCA, HNcoCA, CBCAcoNH and CBCANH spectra were acquired for assignment purposes. All experiments were acquired using time-optimized NMR [52], and were set up using the standard parameter files provided by the software of the instrument provider. Typical acquisition and processing parameters were as above, except that TD for the ^1^H-dimension was 698 and SW was 14.00 ppm. In addition, the TD for ^13^C-dimension was (where applicable) 64 points with linear prediction up to 80. NUS-amount for the 3D experiments was set to 25%. Processed NMR data was imported into CARA [53], for assignment and further analysis. Assigned HSQC spectra for wild type CW in unbound state and bound to corresponding peptides were used to calculate chemical shift perturbations with equation 2 and scaling factor αN of 0.17 [54,55].

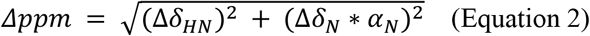

Here, *Δppm* is combined chemical shift; *Δδ*_*HN*_ is amide proton chemical shift, ppm; *Δδ*_*N*_ is nitrogen chemical shift, ppm; *α*_*N*_ is scaling factor.

### Dynamics measurement

Dynamics measurements were performed as described in [24]. Briefly, for determination of heteronuclear NOEs, two ^1^H-^15^N HSQC data sets were recorded. A recycling delay of 3 s was used between transients. Relaxation delays of 20, 60, 80, 100, 200, 400, 600, 800, 1000, 1200, and 1400 ms were recorded for T1 measurements, and relaxation delays of 16, 30, 60, 95, 125, 160, 190, 220, 250, 345, 440, and 500 ms were recorded for T2 measurements. The heteronuclear NOE values were calculated as the ratio of the steady-state intensities measured with and without saturation of the proton magnetization. Data analysis was done in Bruker Dynamics Center 2.5.3 (Bruker BioSpin).

## Acknowledgements

The work was supported through the Norwegian NMR Platform, NNP (grant 226244 / F50) in the form of instruments and experimental time. The authors wish to thank the NNP Bergen Node for facilitating the research. We would also like to thank Carol Issalene for help with protein purification, Jean-Karim Hériché (European Molecular Biology Laboratory) for help with the calculation of tryptophan angles, and Morten Andreas Govasli Larsen for the help with mutagenesis.

## Author contributions

Designed experiments: MB, ØH, ØØF, ØS. Performed experiments: MB, ØH. Analyzed data and interpreted results: MB, OD, ØH, ØØF, RAA, ØS. Wrote drafts: MB, ØH. Revised drafts: MB, ØH, OD, ØS. Prepared Figures: MB, OD, ØH. Data curation: ØH, OD. Formulated research questions and wrote grants: RAA, ØH.

## Abbreviations

PTM: post-translational modification
H3K4meX: histone tail peptide methylated 0-3 times at position Lysine 4
H3K36me3: Histone H3 trimethylation at the Lysine 36
hetNOE: heteronuclear nuclear Overhauser effect
ITC: isothermal titration calorimetry
NMR: nuclear magnetic resonance
NOE: nuclear Overhauser effect
HSQC: heteronuclear single quantum coherence
CSP: chemical shift perturbation

