## Supplementary Information for Binding specificity of ASHH2 CW-domain for "Binding specificity of ASHH2 CW-domain towards H3K4me1 ligand is coupled to its structural stability through its α1-helix" (from another version of this preprint: version 1, 2021-08-12)

Table S1. List of primers.

| Primer[a] | Sequence[b] |
|---|---|
| ASHH2-CW-D870A FW | 5'-TCGATGTGATgccTGCTTTAAATGGCGAC |
| ASHH2-CW-D870A RV | 5'-ACCCAGGCACTCTCTGTG |
| ASHH2-CW-insC866-SFPN-C871 FW | 5'-cctaacTGCTTTAAATGGCGACGAATAC |
| ASHH2-CW-insC866-SFPN-C871 RV | 5'-aaatgaACATCGAACCCAGGCACT |
| ASHH2-CW-I915A FW | 5'-AGAGTTGGGCATAGGACAGGATG |
| ASHH2-CW I915A RV | 5'-TCATTagcTTCTTCATTTGACATCTCTTGAGATTTTGAGC |
| ASHH2-CW L919A FW | 5'-AGAGgctGGCATAGGACAGGATGAAG |
| ASHH2-CW L919A RV | 5'-TCATTAATTTCTTCATTTGACATCTCTTGAG |

[a]FW and RV indicate forward and reverse primers, respectively.
[b]Lower case indicates mutated codons.

Table S2. Histone mimicking peptides.

| Name | Sequence | Manufacturer |
|---|---|---|
| H3 | ARTKQTARY* | LifeTein LLC (95% purity) |
| H3K4me1 | ARTK(me1)QTARY* | LifeTein LLC (95% purity) |
| H3K4me2 | ARTK(me2)QTARY* | LifeTein LLC (95% purity) |
| H3K4me3 | ARTK(me3)QTARY* | LifeTein LLC (95% purity) |

*An additional tyrosine was introduced for concentration determination by absorption at 280 nm (NanoDrop), and an extinction coefficient of 1490 $M^{-1}$ $cm^{-1}$ was used.

Table S3. Summary of ITC data.

| Ligand | Temperature, K | ΔH, kJ/mol | TΔS, kJ/mol | ΔG, kJ/mol | $K_d$, μM |
|---|---|---|---|---|---|
| CW wild type | | | | | |
| H3K4me1 | 298.15 | -89.25±8.58 | -57.37±7.14 | -31.88±2.78 | 1.31±0.32 |
| H3K4me2 | 298.15 | -83.64±3.11 | -53.18±3.23 | -30.46±0.15 | 4.61±0.28 |
| H3K4me3 | 298.15 | -60.14±1.38 | -32.47±1.49 | -27.67±0.13 | 14.18±0.74 |
| D870A | | | | | |
| H3K4me1 | 298.15 | -79.22±4.81 | -45.70±4.10 | -33.52±0.71 | 1.37±0.32 |
| H3K4me2 | 298.15 | -93.33±3.65 | -64.03±3.59 | -29.29±0.23 | 7.48±0.72 |
| H3K4me3 | 298.15 | -57.47±2.37 | -30.45±2.65 | -27.02±0.29 | 18.53±2.17 |
| L919A | | | | | |
| H3K4me1 | 298.15 | -76.92±2.31 | -48.84±1.97 | -28.08±0.42 | 12.17±2.10 |

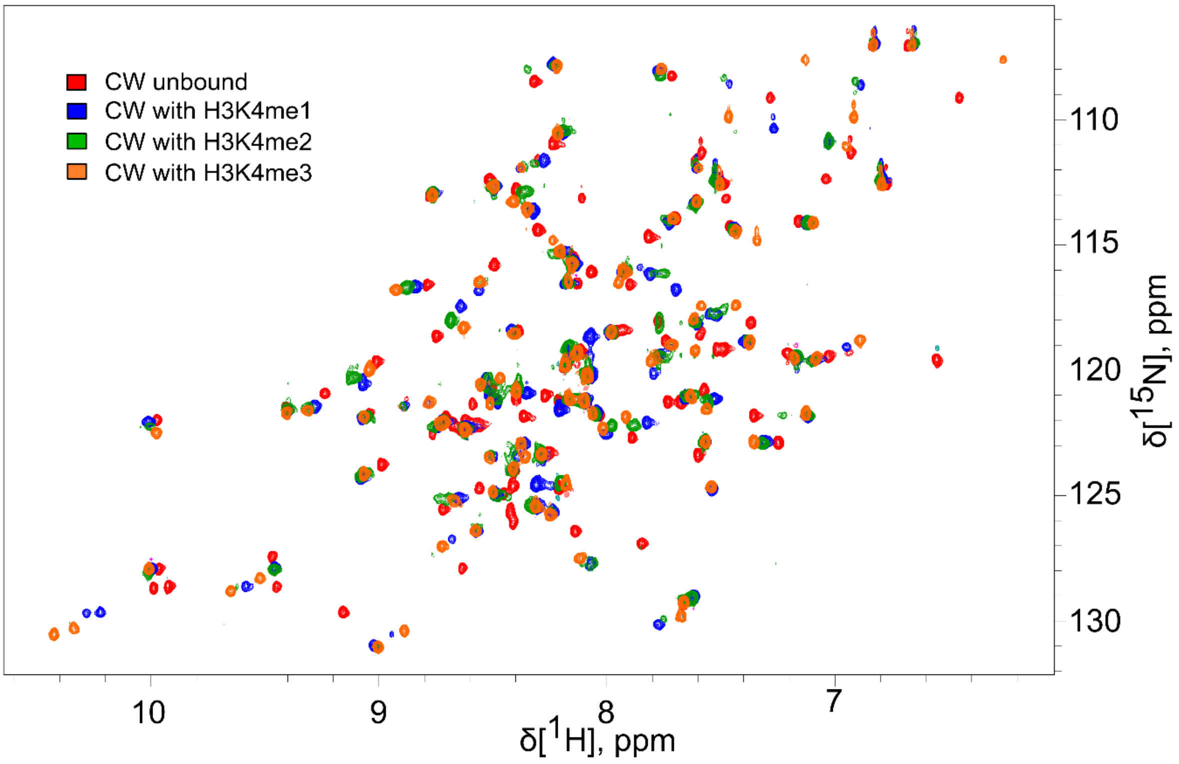


### Figure S1.

Overlaid HSQC spectra of CW wild type in free state (red) and bound to H3K4me1 (blue), H3K4me2 (green) and H3K4me3 (orange) peptides.

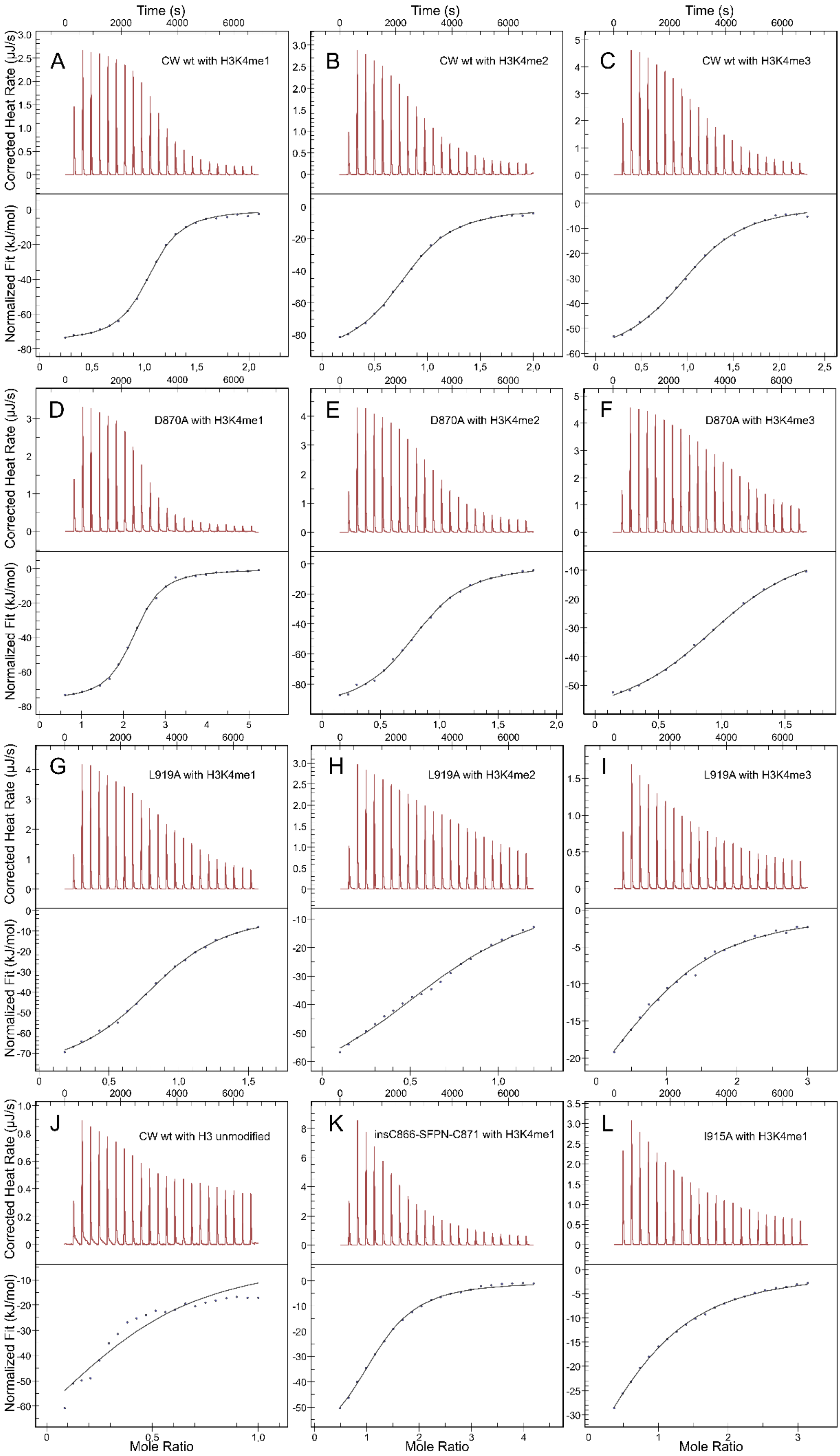
Time (s)
Corrected Heat Rate (μJ/s)
Normalized Fit (kJ/mol)
A
CW wt with H3K4me1
B
CW wt with H3K4me2
C
CW wt with H3K4me3
D
D870A with H3K4me1
E
D870A with H3K4me2
F
D870A with H3K4me3
G
L919A with H3K4me1
H
L919A with H3K4me2
I
L919A with H3K4me3
J
CW wt with H3 unmodified
K
insC866-SFPN-C871 with H3K4me1
L
I915A with H3K4me1
Mole Ratio

### Figure S2

Titration curves and integrated data of: A-C – CW wt with H3K4me1/2/3 peptides; D-F – D870A with H3K4me1/2/3 peptides; G-I – L919A with H3K4me1/2/3 peptides; J – CW wt with H3 unmodified peptide; K – insC866-SFPN-C871 with H3K4me1 peptide; L – I915A with H3K4me1 peptide. In H-L no sigmoidal curve could be fitted.
